# NK cells promote cancer immunoediting through tumor-intrinsic loss of interferon-stimulated gene expression

**DOI:** 10.1101/2020.11.23.394353

**Authors:** Yung Yu Wong, Luke Riggan, Edgar Perez-Reyes, Christopher Huerta, Matt Moldenhauer, Timothy E. O’Sullivan

## Abstract

Natural killer (NK) cells are innate lymphocytes that constantly patrol host tissues against transformed cells in a process known as cancer immunosurveillance. Previous evidence in mice has demonstrated that NK cell-derived IFN-γ can promote immunoevasion by sculpting the immunogenicity of developing tumors in a process known as cancer immunoediting. This process involves the elimination of highly immunogenic “unedited” tumor cells followed by the eventual escape of less immunogenic “edited” tumor cell variants that are able to escape recognition or elimination by the immune system. Here, we show that NK cell-edited fibrosarcomas decrease the expression of 17 conserved IFN-γ-inducible genes compared to unedited tumor cells. High expression of 3 of these identified genes (*Psmb8, Trim21, Parp12*) in human tumor samples correlates with enhanced survival in breast cancer, melanoma, and sarcoma patients. While NK cell-edited fibrosarcomas displayed resistance to IFN-γ growth suppression *in vitro*, functional knockouts of individual interferon stimulated genes (ISGs) were not required for outgrowth of unedited tumor cell lines *in vitro* and *in vivo* compared to complete loss of IFN signaling. Furthermore, knockout of IFN-γ-intrinsic signaling via deletion of *Ifngr* in edited B16 F10 and E0771 LMB metastatic cancer cell lines did not impact host survival following lung metastasis. Together, these results suggest that unedited tumors can be selected for decreased IFN-γ signaling to evade immune responses *in vivo*, and as a consequence, tumor-extrinsic IFN signaling may be more important for potentiating durable anti-tumor responses to advanced solid tumors.

## Introduction

Research from the past decade has demonstrated the exciting potential of harnessing a patient’s immune system to prevent or delay tumor progression and metastasis^1^. Although immunotherapies targeting adaptive lymphocytes (i.e. antigen-specific T cells) have proven successful in certain circumstances, most cancer patients ultimately do not respond and can suffer complications due to unrestrained immune responses^1, 2^. Recent studies in mice and humans have revealed that tumor-intrinsic mutations in the interferon (IFN)-γ receptor signaling pathway correlate with poor outcomes in response to checkpoint blockade immunotherapy^3, 4^, suggesting that resistance to IFN-γ signaling is a clinically relevant mechanism of tumor immunoevasion. Thus, there is an increasing need to understand the mechanisms by which IFN-γ-inducible genes suppress tumor progression in order to develop more effective therapeutic strategies for the large proportion of cancer patients that may never respond to current immunotherapies.

Natural killer (NK) cells are innate lymphocytes that constantly patrol host tissues against transformed cells in a process known as cancer immunosurveillance. Previous evidence in mice has demonstrated that NK cell-derived IFN-γcan promote immunoevasion by sculpting the immunogenicity of developing tumors in a process known as cancer immunoediting^5, 6^. This process involves the elimination of highly immunogenic “unedited” tumor cells followed by the eventual escape of less immunogenic “edited” tumor cell variants that are able to escape recognition or elimination by the immune system^5, 6^. Here, we show that NK cell-edited fibrosarcomas decrease the expression of 17 conserved IFN-γ-inducible genes compared to unedited tumor cells. High expression of 3 of these identified genes (*Psmb8, Trim21, Parp12)* in human tumor samples correlates with enhanced survival in breast cancer, melanoma, and sarcoma patients. While NK cell-edited fibrosarcomas displayed resistance to IFN-γ growth suppression *in vitro*, functional knockouts of individual interferon stimulated genes (ISGs) were not required for outgrowth of unedited tumor cell lines *in vitro* and *in vivo* compared to complete loss of IFN signaling. Furthermore, knockout of IFN-γ-intrinsic signaling via deletion of *Ifngr* in edited B16 F10 and E0771 LMB metastatic cancer cell lines did not impact host survival following lung metastasis. Together, these results suggest that unedited tumors can be selected for decreased IFN-γ signaling to evade immune responses *in vivo*, and as a consequence, tumor-extrinsic IFN signaling may be more important for potentiating durable anti-tumor responses to advanced solid tumors.

## Results

Previous studies have shown that MCA-induced fibrosarcomas developed in Rag2^-/-^ or Rag2^-/-^ x γc^-/-^ mice develop more tumors that are recognized and rejected by the immune system of WT mice^5, 6^. “Unedited” fibrosarcomas derived from *Rag2*^-/-^ x *Il2rg*^-/-^ mice can be transplanted in *Rag2*^-/-^ mice and harvested to produce daughter “edited” tumor cell lines that progressively grow in WT mice. While NK cells and IFN-γ were found to be required for this editing process *in vivo*, the tumor-intrinsic changes in gene expression that occur during NK cell-mediated immunoediting remained unknown. To determine this, we transplanted the unedited fibrosarcoma line 7357dsRed into NK cell sufficient (*Rag2*^-/-^) or NK cell deficient (*Rag2*^-/-^ x *Il2rg*^-/-^) mice to generate NK cell-passaged (NP) and control-passaged (CP) daughter tumor cells respectively (**Fig. 1A**). RNAseq analysis of *ex vivo* sorted dsRed^+^ tumor cells revealed higher transcriptional similarity between NP rather than CP tumor cells, suggesting NK-cell dependent influence on the tumor transcriptome *in vivo* (**Fig. 1B**). Interestingly, analysis of the differentially expressed genes (DEGs) between CP and NP tumor cells revealed a dramatic loss of type II interferon stimulated gene (ISG) expression, with an increase in a small subset of genes (**Fig. 1C,D**). To determine the relevance of these findings, we compared the list of DEGs down in NP tumors to a previously validated dataset of mouse and human ISGs^7^. Of the 164 genes downregulated in NP tumors, 64 genes were found to be ISGs, with 17 expressed conserved in mouse and human fibroblasts (**Fig. 2A**). Using publicly available datasets from the Cancer Genome Atlas (TCGA), we determined that 13 of these genes predicted better survival outcomes for SARC (sarcoma) and SKCM (melanoma) tumors (**data not shown**), and 3 genes (*TRIM21, PSMB8, PARP12*) predicted better survival outcomes for BRCA (breast), SKCM, and SARC tumors (**Fig. 2B,C**).

**Figure 1.**
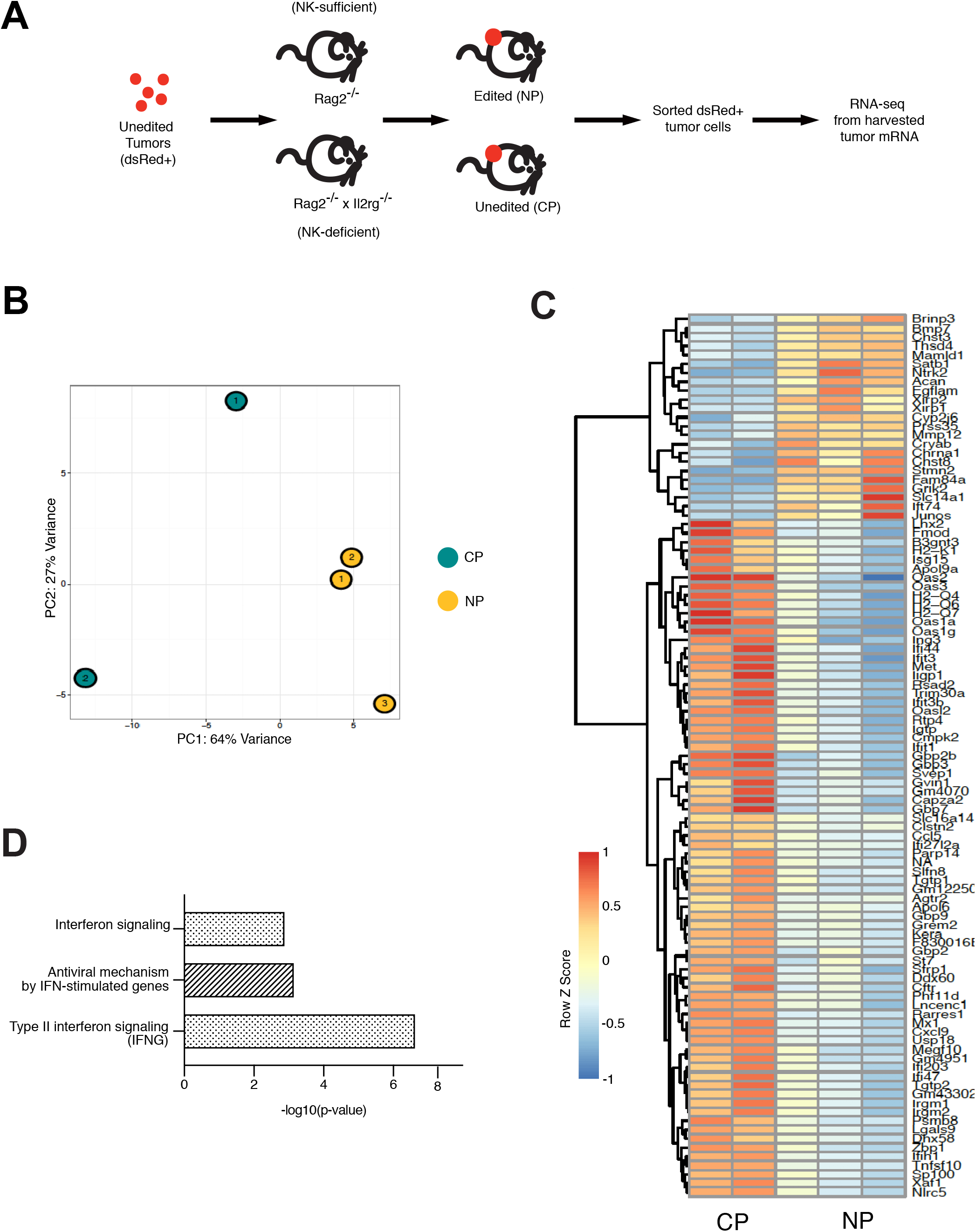
NK cell dependent immunoediting reveals tumor-intrinsic suppression of IFN-γ-inducible genes. The MCA-induced fibrosarcoma cell line 7357 derived from *Rag2*^-/-^ x *Il2rg*^-/-^ mice was transduced with lentivirus and selected to stably express dsRED. dsRED^+^ tumor cells were transplanted into either NK cell sufficient *Rag2*^-/-^ or NK cell deficient *Rag2*^-/-^ x *Il2rg*^-/-^ mice. After 15 days, tumors were harvested into single cell suspensions. dsRED^+^ tumor cells were sorted by flow cytometry and mRNA was harvested to generate libraries for RNA-sequencing. (**A**) Schematic of experiment (**B**) Principle component analysis of the transcriptome of *Rag2*^-/-^ x *Il2rg*^-/^ passaged 7357 tumor cells (Control Passaged, CP) compared to *Rag2*^-/-^ passaged 7357 tumor cells (NK cell passaged, NP) (**C**) Heat map displays differentially expressed genes (DEGs) between NP and CP tumor cells. **(D)** Gene enrichment analysis of DEGs between NP and CP tumor cells.

**Figure 2.**
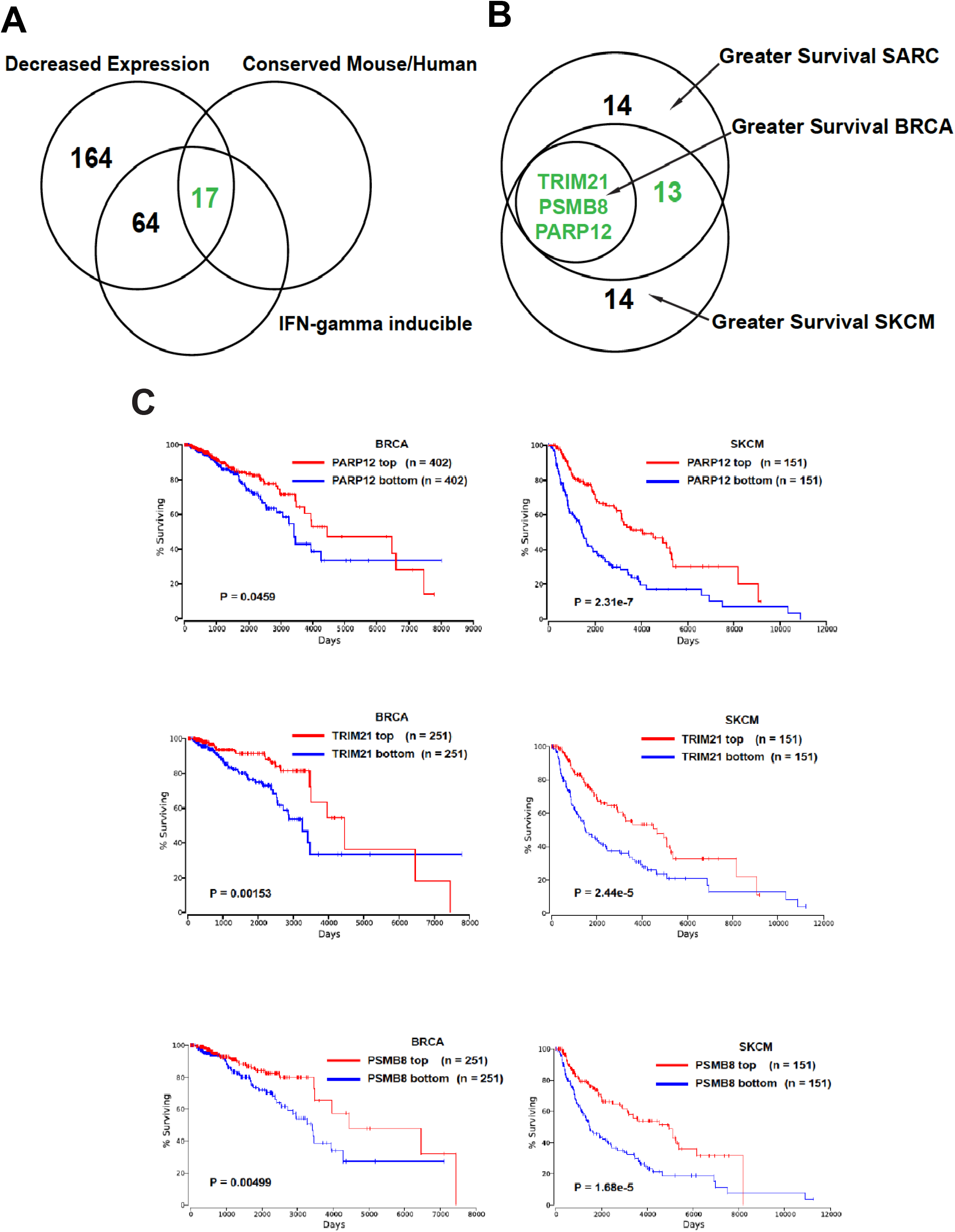
ISGs conserved between human and mouse correlate with patient survival. (**A**) Gene overlap between downregulation in NP vs CP tumors, IFN-γ-inducible genes in mouse fibroblasts from interferome datasets, and IFN-γ-inducible genes in human fibroblasts from interferome datasets (**B**) Prognostic value of conserved IFN-gamma-inducible genes identified for overall breast (BRCA), sarcoma (SARC), and skin (SKM) cancer patients comparing top and bottom quartiles of expression. (**C**) Prognostic value of *Parp12, Psmb8*, and *Trim21* expression levels for indicated cancer patient overall survival comparing top and bottom quartiles of expression.

Because NP tumors displayed reduced ISG expression directly *ex vivo*, we next tested whether these cells were resistant to IFN-γ signaling While CP tumors displayed a 20% decrease in proliferation compared to untreated controls, IFN-γ treated NP tumors proliferated to the same extent as untreated NP and *Stat1*^-/-^ CP tumor cells (**Fig. 3A**), suggesting that NK cell editing *in vivo* can lead to tumor escape of IFN-γ-mediated growth restriction *in vitro*. This process likely occurs *in vivo* through increased fitness of fibrosarcoma clones with indels in multiple ISGs, because CRISPR-mediated knockout of these proteins did not display resistance to IFN-γ-mediated growth restriction *in vitro* or outgrowth in WT mice (**Fig. 3B,C**) and CRISPR-induced knockout indels of individual ISGs downregulated in NP tumors were not selected during culture with IFN-γ (**Fig. 3D**). In support of this hypothesis, functional *Stat1* knockout indels in CP tumor cells showed enrichment compared to WT *Stat1* sequences following culture with IFN-γ, and *Stat1*^-/-^ CP tumors progressively grew following transplantation in WT mice (**Fig. 3D,E**).

**Figure 3.**
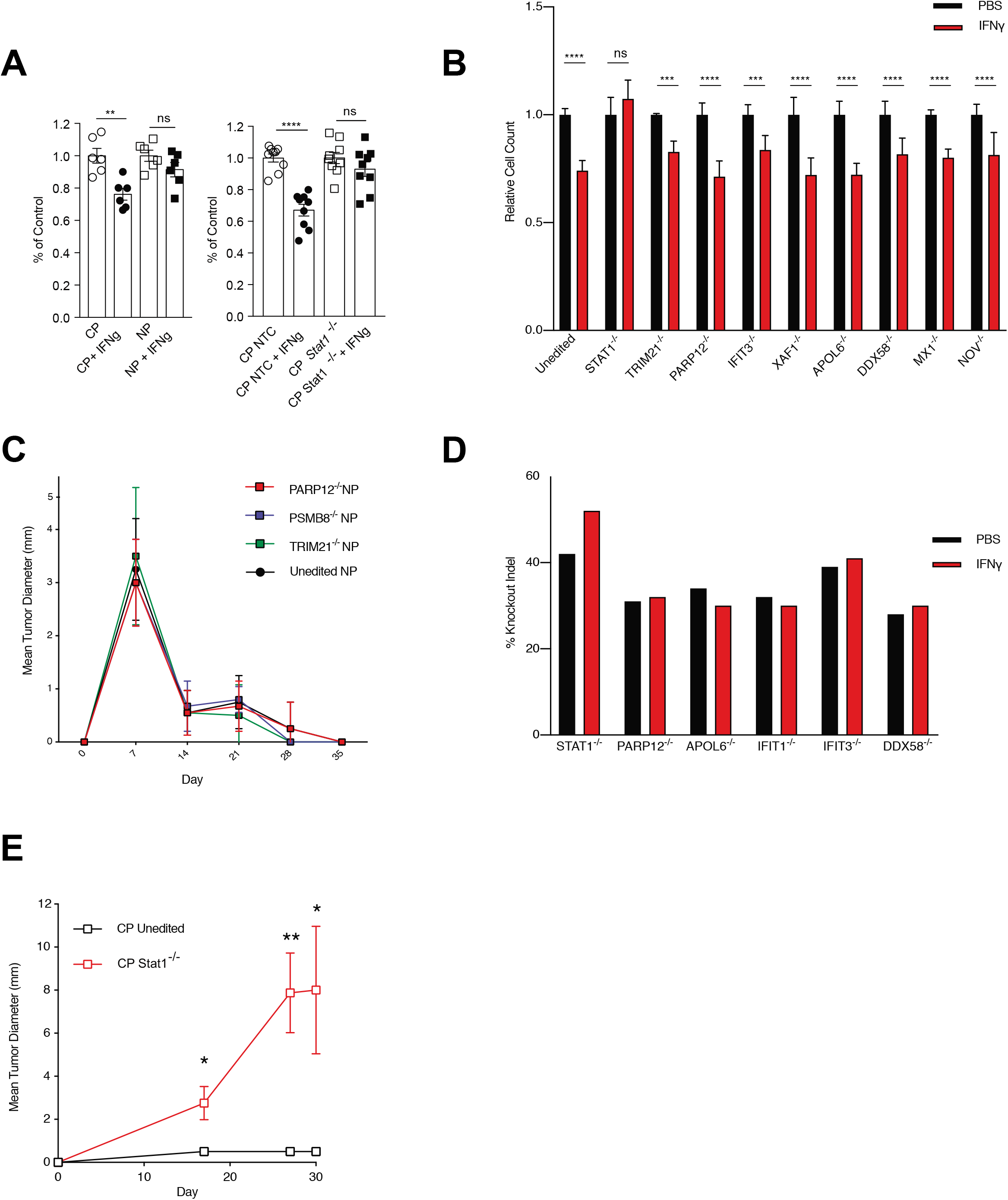
Individual ISG KO tumor cell lines are not resistant to IFN-mediated growth suppression *in vitro* or *in vivo*. **(A)** CP, NP, CP Cas9 non-targeting control (NTC), and CP Stat1 KO 7357 fibrosarcoma tumor cells were grown in the presence or absence of 15 ng/ml IFN-γ for 4 days. Viable tumor cells were counted by flow cytometry. Results are shown as the percentage of non-IFN-γ treated controls for each cell line. **(B)** Tumor cell lines were cultured in either 15ng/mL IFN-γ or PBS (control) for 4 days. Cell counts were determined by fluorescence-activated cell sorting and normalized. **(C)** Tumor cell lines were injected into the right flank of wild type C57BL/6J mice via subcutaneous injection. Unedited tumors were injected on the left flank as control. Mean tumor diameters were measured every week for 5 weeks. **(D)** Tumor cell lines were cultured in 15ng/mL IFN-γ for 4 days and the selection for KO cells was determined by Inference of CRISPR Editing (ICE) analysis. **(E)** Single cell STAT1 KO clones and unedited tumors were injected into wild type mice and the mean tumor diameters were measured every 10 days for 30 days. Samples were compared using two-tailed Student’s t test with Welch’s correction, assuming unequal SD, and data are presented as individual points with the mean ± SEM (*p<0.05, **p<0.01, ***p<0.001, ****p<0.0001). Linear regression and Pearson Correlation analysis with 95% confidence intervals were conducted. p < 0.05 was considered significant.

While CP tumors are useful for studying initial immune responses to developing cancer *in vivo*, established mouse tumor models are derived from mice with an intact immune system, and therefore have already been edited. To test whether IFN-γ signaling was required for tumor-intrinsic growth suppression in other edited tumor types, we generated *Ifngr*^-/-^ B16-F10 metastatic melanoma and E0771 LMB metastatic breast cancer cell lines that have been shown to be IFN-γ-sensitive *in vivo* previously^8, 9^. Interestingly, we did not observe a difference in the survival of mice receiving WT or *Ifngr*^-/-^ B16-F10 or E0771 LMB metastatic tumor cells *i.v*. (**Fig 4A,B**) These results suggest that tumor-extrinsic IFN-γ signaling may be necessary for immune-mediated control of edited cancers *in vivo*, likely as a consequence of immunoediting of tumor-intrinsic IFN-γ signaling early during tumor development.

**Figure 4.**
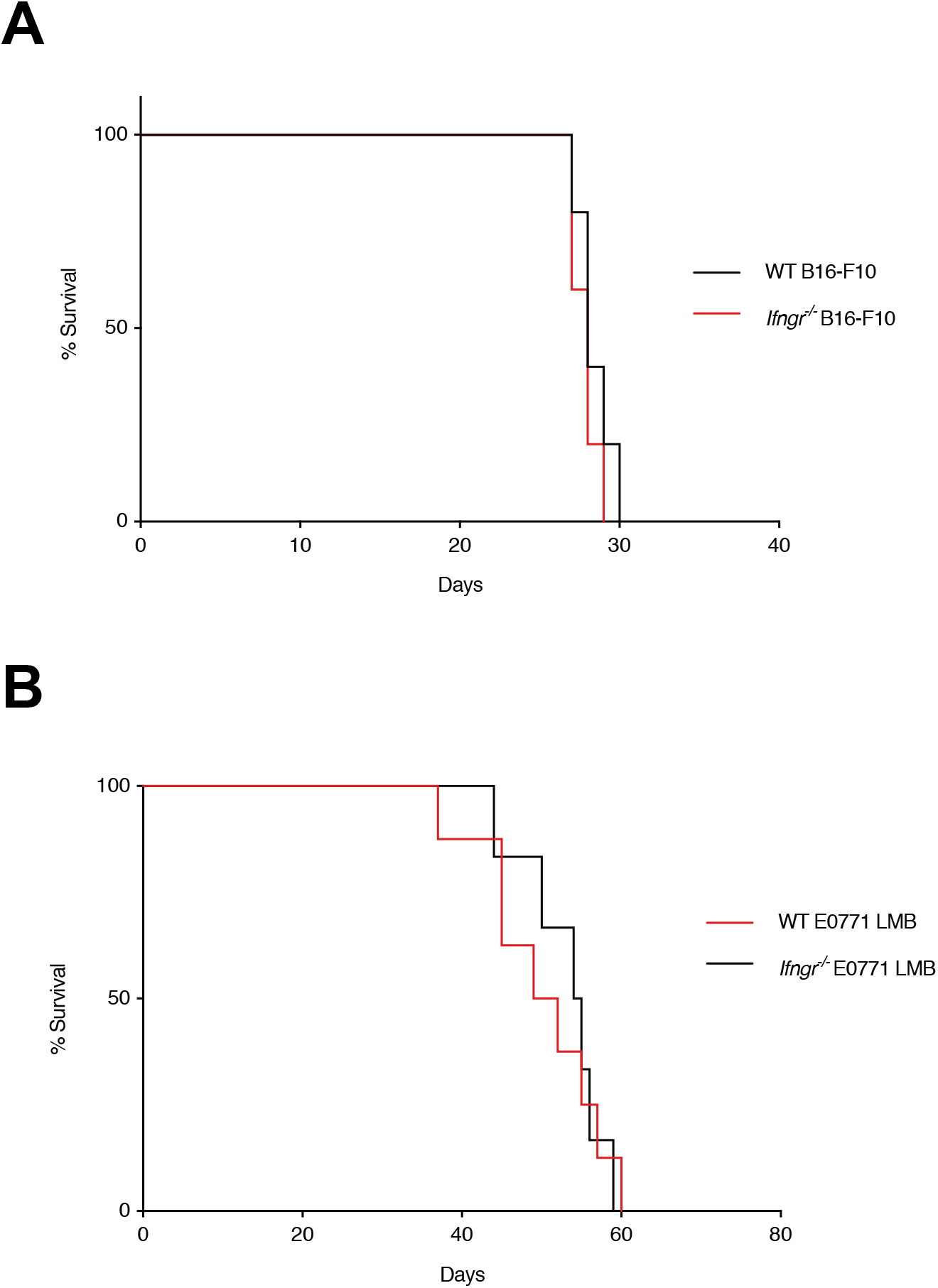
Edited tumor cell lines do not require intrinsic IFN-γ signaling for metastasis *in vivo*. **(A) (B)** Unedited or *Ifngr*^-/-^ B16-F10 and E0771 LMB cells were injected into wild type C57BL/6J mice via intravenous injection. Percent survival was determined over 60 days.

## Discussion

IFN-γ production and activation of NK cells in the tumor microenvironment correlates with better survival outcomes in human cancer^10, 11^. Similarly, previous studies have shown that mice lacking STAT1 are more susceptible to carcinogen MCA-induced tumors^5^, further consolidating the role of IFN-γ signaling in tumor suppression. However, our results suggest that NK cell derived IFN-γ may act as a double-edged sword as it can also facilitate immunoevasion of the less immunogenic tumors via immunoediting. Our data supports the hypothesis that tumor-intrinsic suppression of the IFN-γ signaling pathway is key to tumor immunoevasion of immunogenic tumor cells during cancer development.

While previous *in vivo* CRISPR screens found that individual tumor-expressed ISGs are important in T cell dependent antitumor responses^3^, checkpoint blockade nonresponders show multiple deletions in ISGs rather than single gene deletions^4^. These results suggest that immune evasion in tumor cells is likely to be more complex *in vivo*, and multiple ISG deletions may be necessary to yield significant immunoevasion. Our results support this hypothesis by showing the redundant nature of individual ISGs in the rejection of highly immunogenic tumors *in vivo*. The contribution of individual ISGs is likely complex due to a redundancy in their roles in antigen presentation and cell cycle arrest^12^. Teasing out the specific combinations of ISG gene deletions that render tumor insensitive to lymphocyte-based therapies will be needed in future studies. While our data only investigated the roles of ISGs in tumors, recent studies indicate that the expression and function of ISGs are likely to be cell type specific. ISGs differentially expressed in tumors and immune cells could have opposing functions, counteracting each other in the context of lymphocyte activation in the tumor microenvironment^13^. Our data aligns with previous findings that tumor-extrinsic IFN signaling may be more important for anti-tumor responses in advanced tumors.

## Methods

### RNA sequencing

For RNA sequencing, RNA was isolated from sorted cell populations from WT mice using TRIzol (Invitrogen) followed by SMARTer amplification and Illumina next-generation sequencing. RNA and ATAC sequencing reads were aligned and pre-processed as described previously^14^. RNA sequencing reads were aligned using STAR (2.5.3a-foss-2016b) with filters --outFilterMultimapNmax 1 --outFilterMismatchNmax 999 -- outFilterMismatchNoverLmax 0.02 --alignIntronMin 20 --alignIntronMax 1000000 -- alignMatesGapMax 1000000 to allow for stringent alignment of unique reads to the mouse (mm10) genome^15^. Counts were made using BEDtools 2.27.1 coveragebed function. Raw count files were processed using DESeq2, removing genes with less than 50 counts.

### Selection of guide RNAs

Webtool CHOPCHOPv2^16^ was used to select target sites for CRISPR/Cas9. Single guide RNAs (sgRNAs) within the first three exons were selected to minimize the number of off target sites. MIT CRISPR design database^17^ was used to cross-reference sgRNA selection by evaluating the ranking of target sites according to the MIT database’s algorithm.

### Production of lentivirus-based CRISPR-Cas9 knockout tumor cell lines

According to the LentiCRISPRv2 protocol^18^, CRISPR/Cas9 LentiCrisprV2 plasmid with sgRNAs were generated via cloning and transformation of Stbl3 E.Coli. Transfection reagents pSPAX and pMDG.2 were generated by ampicillin-resistant Stbl3 E.Coli with the corresponding plasmid. Briefly, the bacteria were grown in 100μg/mL sterile ampicillin LB solution in a shaking incubator at 37°C for 6-8hours. 200uL of the culture was placed in 25mL of sterile ampicillin LB solution and incubated in a shaking incubator at 37°C for 12-18 hours. Bacterial DNA was isolated using QIAGEN Plasmid MidiPrep Kit and suspended in DEPC-treated water. Next, human embryonic kidney cells (HEK) 293T were grown in a T75 flask with Dulbecco’s modified eagle medium (DMEM) (Gibco) + 10% fetal bovine serum (FBS) at 37°C and plated onto a six-well plate after reaching 75%-85% confluency. After reaching 75%-85% confluence on the six-well plate, the 293T cells were transfected with the previously isolated plasmid DNA as following: mixture 1 contains 2.5μg Midiprep-isolated plasmid DNA, 1.9μg pSPAX, 1.25μg pMDG.2 and 11μl Lipofectamine P3000 transfection reagent in 500μl OPTI-MEM. This mixture was combined with mixture 2: 500μl OPTI-MEM and 11μl Lipofectamine solution. 330μl of the combined mixture was added to the plated 293T (3 wells per sgRNA) and the plate was incubated for 5 hours at 37°C, followed by replacement of media with DMEM + 10% FBS. 7357 fibrosarcoma cells were grown in CR-10 media (RPMI 1640 Gibco) and seeded onto a six-well plate at the density of 30,000 cells/ well the same day as the initial transfection. 48 hours after initial transfection, supernatant from 293T cells was harvested and syringe filtered through 0.45μm filter and placed on 7357 fibrosarcoma cells (3mL total per sgRNA per well). The following day, 7357 tumor cells were trypsinized with 0.25% trypsin-EDTA and expanded into a T75 flask with CR-10 media. 48 hours later, fresh CR-10 media was added to the T75 alongside blasticidin at the concentration of 10μg/mL. After reaching 70% confluence, the edited cell lines were expanded into T150 flasks in order to be frozen down or extract genomic DNA with Zymo Research Quick-DNA kit. Inference of CRISPR Editing (ICE) analysis^19^ was conducted on the genomic DNA to characterize mutations in KO cell lines and evaluate the editing efficiency.

### In vitro tumor growth assay

Tumor cell lines were grown in T150 flask with 20mL CR-10 media until 70-80% confluence. Cells were plated at concentration of 20,000 cells per well with 1mL serum starved CR-10, and 3mL were added after. 20 hours later, serum starved CR-10 media was replaced with 3mL CR-10, IFN-γ at 15 ng/mL and blasticidin at 10μg/mL. Four days later, 7357 cells were washed with 1mL Hank’s balanced salt solution (HBSS), harvested and suspended in 1mL PBS. Fluorescence-activated cell sorting was used to count cells.

### Determining knockout indel selection by IFN-γ

CRISPR edited tumor cell lines were grown in IFN-γ growth assays harvested from the previously described IFN-gamma growth assays and the genomic DNA was extracted using Zymo Research Quick-DNA kit. ICE analysis was conducted on the genomic DNA to determine the knockout indel percentage.

### In vivo tumor growth

Tumor cell lines were grown to confluence in T150 flask. 24 hours before injection, cells were split into new T150 at a 1:3 ratio. The following day cells were harvested and spun down at 1500 RPM for 10 minutes at 4°C. Supernatant was decanted and 5mL HBSS was added, pellet was resuspended into single cell solution and spun down again at 1500 RPM for 3 minutes at 4°C. Supernatant was decanted and pellet was resuspended to make a final solution of 1 million cells per 200μl. Tumor cells were then delivered via subcutaneous injection to wild type C57BL/6J mice at 1 million tumor cells per mouse. The 7357 KO line was delivered into the right flank and the unedited control into the left flank of each mouse. From Day 0 to 35, tumor sizes were measured every week.

## Author Contributions

Y.Y.W., L.R., and T.E.O. designed the study; Y.Y.W., E.P.R, C.H., M.M., L.R., and T.E.O performed the experiments; T.E.O. performed RNA-seq bioinformatics analysis; Y.Y.W. and T.E.O. wrote the manuscript.

## References

1. Sharma, P., Hu-Lieskovan, S., Wargo, J.A. & Ribas, A. Primary, Adaptive, and Acquired Resistance to Cancer Immunotherapy. Cell 168, 707–723 (2017).

2. Michot, J.M. et al. Immune-related adverse events with immune checkpoint blockade: a comprehensive review. Eur J Cancer 54, 139–148 (2016).

3. Gao, J. et al. Loss of IFN-gamma Pathway Genes in Tumor Cells as a Mechanism of Resistance to Anti-CTLA-4 Therapy. Cell 167, 397–404 e399 (2016).

4. Zaretsky, J.M. et al. Mutations Associated with Acquired Resistance to PD-1 Blockade in Melanoma. N Engl J Med 375, 819–829 (2016).

5. Schreiber, R.D., Old, L.J. & Smyth, M.J. Cancer immunoediting: integrating immunity’s roles in cancer suppression and promotion. Science 331, 1565–1570 (2011).

6. O’Sullivan, T. et al. Cancer immunoediting by the innate immune system in the absence of adaptive immunity. J Exp Med 209, 1869–1882 (2012).

7. Rusinova, I. et al. Interferome v2.0: an updated database of annotated interferon-regulated genes. Nucleic Acids Res 41, D1040–1046 (2013).

8. Benci, J. L., Xu, B., Qiu, Y., Wu, T. J., Dada, H., Twyman-Saint Victor, C., Cucolo, L., Lee, D., Pauken, K. E., Huang, A. C., Gangadhar, T. C., Amaravadi, R. K., Schuchter, L. M., Feldman, M. D., Ishwaran, H., Vonderheide, R. H., Maity, A., Wherry, E. J., & Minn, A. J. (2016). Tumor Interferon Signaling Regulates a Multigenic Resistance Program to Immune Checkpoint Blockade. Cell, 167(6), 1540–1554.e12.

9. Kearney, C. J., Vervoort, S. J., Hogg, S. J., Ramsbottom, K. M., Freeman, A. J., Lalaoui, N., … Oliaro, J. (2018). Tumor immune evasion arises through loss of TNF sensitivity. Science Immunology, 3(23).

10. Boudreau JE, Hsu KC. Natural Killer Cell Education and the Response to Infection and Cancer Therapy: Stay Tuned. Trends Immunol. 2018; 39(3):222–239.

11. Zhang, S., Liu, W., Hu, B., Wang, P., Lv, X., Chen, S., Shao, Z., Prognostic Significance of Tumor-Infiltrating Natural Killer Cells in Solid Tumors: A Systematic Review and Meta-Analysis. Frontiers in Immunology 11, 1242 (2020).

12. de Veer, M.J., Holko, M., Frevel, M., Walker, E., Der, S., Paranjape, J.M., Silverman, R.H. and Williams, B.R.G. (2001), Functional classification of interferon-stimulated genes identified using microarrays. Journal of Leukocyte Biology, 69: 912–920.

13. Benci J.L., Johnson L.R., Choa R., Xu Y., Qiu J., Zhou Z., Xu B., Ye D., Nathanson K.L., June C.H., Wherry E.J., Zhang N.R, Ishwaran H., Hellmann M.D., Wolchok J.D., Kambayashi T., Minn A.J.. Opposing Functions of Interferon Coordinate Adaptive and Innate Immune Responses to Cancer Immune Checkpoint Blockade. Cell. 2019 Aug 8;178(4):933–948.e14.

14. Weizman OE, Adams NM, Schuster IS, Krishna C, Pritykin Y, Lau C, et al. ILC1 Confer Early Host Protection at Initial Sites of Viral Infection. Cell 2017, 171(4): 795–808 e712.

15. Tokuyama M, Kong Y, Song E, Jayewickreme T, Kang I, Iwasaki A. ERVmap analysis reveals genome-wide transcription of human endogenous retroviruses. Proc Natl Acad Sci U S A 2018, 115(50): 12565–12572.

16. Labun, K., Montague, T. G., Gagnon, J. A., Thyme, S. B., & Valen, E. (2016). CHOPCHOP v2: A web tool for the next generation of CRISPR genome engineering. Nucleic Acids Research, 44(W1).

17. Hsu, P., Scott, D., Weinstein, J. et al. DNA targeting specificity of RNA-guided Cas9 nucleases. Nat Biotechnol 31, 827–832 (2013).

18. Sanjana, N.E., Shalem, O. & Zhang, F. Improved vectors and genome-wide libraries for CRISPR screening. Nat Methods 11, 783–784 (2014).

19. Brinkman, E.K., Chen, T., Amendola, M. & van Steensel, B. Easy quantitative assessment of genome editing by sequence trace decomposition. Nucleic Acids Res 42, e168 (2014).

